# Metacommunity framework and its core terms entanglement

**DOI:** 10.1101/2021.09.26.461405

**Authors:** Jurek Kolasa, Matthew P. Hammond, Joyce Yan

**Author notes:** Correspondence: Department of Biology, McMaster University, Hamilton, ON, L8S 4K1, Canada.

## Abstract

The metacommunity framework links space and ecological processes but is vulnerable to complex entanglement among its integral components. Most ecological processes are context-dependent. However, when ecological theories show it, they may be seriously crippled unless they explicitly tackle it. Otherwise, findings emerging from accumulated cases will be of limited value and likely remain ambiguous or misleading. Specifically, interactions among the core terms of metacommunity theory interact in complex ways that we identify as entanglement. We employ four core dimensions to alleviate this issue and create a space where various studies converse and effectively complement each other irrespective of the case specifics. The dimensions encompass the metacommunity empirical domain: (1) inter-habitat differences, (2) species habitat specialization, (3) effective dispersal, and (4) species interactions (negative to positive). Then, we assess the entanglement effects by testing that (a) ***changing values in one dimension, with others constant, alters study conclusions***, and (b) ***these effects increase and dominate when integral dimensions interact reciprocally***. As a metric, we analyzed species diversity in a stochastic, agent-based, unified metacommunity model, UMM, where species move, select habitats, reproduce, and interact. In the simulations, each dimension has four or five levels spanning a broad spectrum of conditions. The exercise strongly supports both hypotheses. It also suggests that positive interactions, in contrast to the popular emphasis, promote biodiversity more than negative ones like competition or predation. The proposed integrated conceptual system can expand to include meta-ecosystems, habitat gradients, and other processes. Thus, it can offer a unified approach to spatial processes in ecology. Finally, by combining the four dimensions into one interactive system, we identify a rich array of lower-level hypotheses that inevitably emerge from this system. The hypotheses’ shared origin anchors individual studies in coherent structure to advance sound generalizations.

## Introduction

The outcomes of ecological processes are often context-dependent (1). For example, the growth of a population depends on the species’ life history, resources, other species, environment, or population size. Such context-dependency, CD, reduces success in searching for generalizations and testing hypotheses (2) in natural settings. Not surprisingly, multispecies processes are not free of CD. A severe challenge arises when the integral building blocks of a theory are also context-dependent, as appears to be the case with the metacommunity framework, MCF. This unique and consequential case deserves a term: conceptual entanglement, or CE.

What may appear to be an obstacle also presents an opportunity. The MCF possesses promising strengths for developing ecological theory appropriate for multispecies systems. Its scope includes the ecology of discrete and continuous processes in space where species interactions, dispersal, diversity patterns, ecosystem processes, and local/regional scaling define its primary foci and domain (3). The MCF appeal arises from many testable predictions for a broad range of empirical processes. This broad ability led to MCF’s exponential growth (Web of Science: from 3 papers in 2000 to 187 in 2021) and significant monographic treatments (4, 5). Yet, to gain power, MCF must be more precise about its foundations. Furthermore, the entanglement of the core terms implies the existence of complex relationships that require articulation and testing.

Its rapid expansion highlights two areas of concern, potentially impeding the consistency and efficiency of MCF. The first is more evident than the second: MCF accrued diverse approaches (2, 4, 5), ad hoc components, modifications, foundational ambiguities, and gaps typical during a theory’s rapid growth. Metacommunity models with inconsistent core dimensions, the role of competition-dispersal tradeoffs, space considerations, and others (Table 1) aim to satisfy calls (e.g., (6)) for integrating existing theoretical approaches. The proposed solutions lack a clear inference space to differentiate between the ideas and give context to the results (7). Emerging topics, like food webs (e.g., (7, 8)), add to the sense of challenge.

**Table 1.**
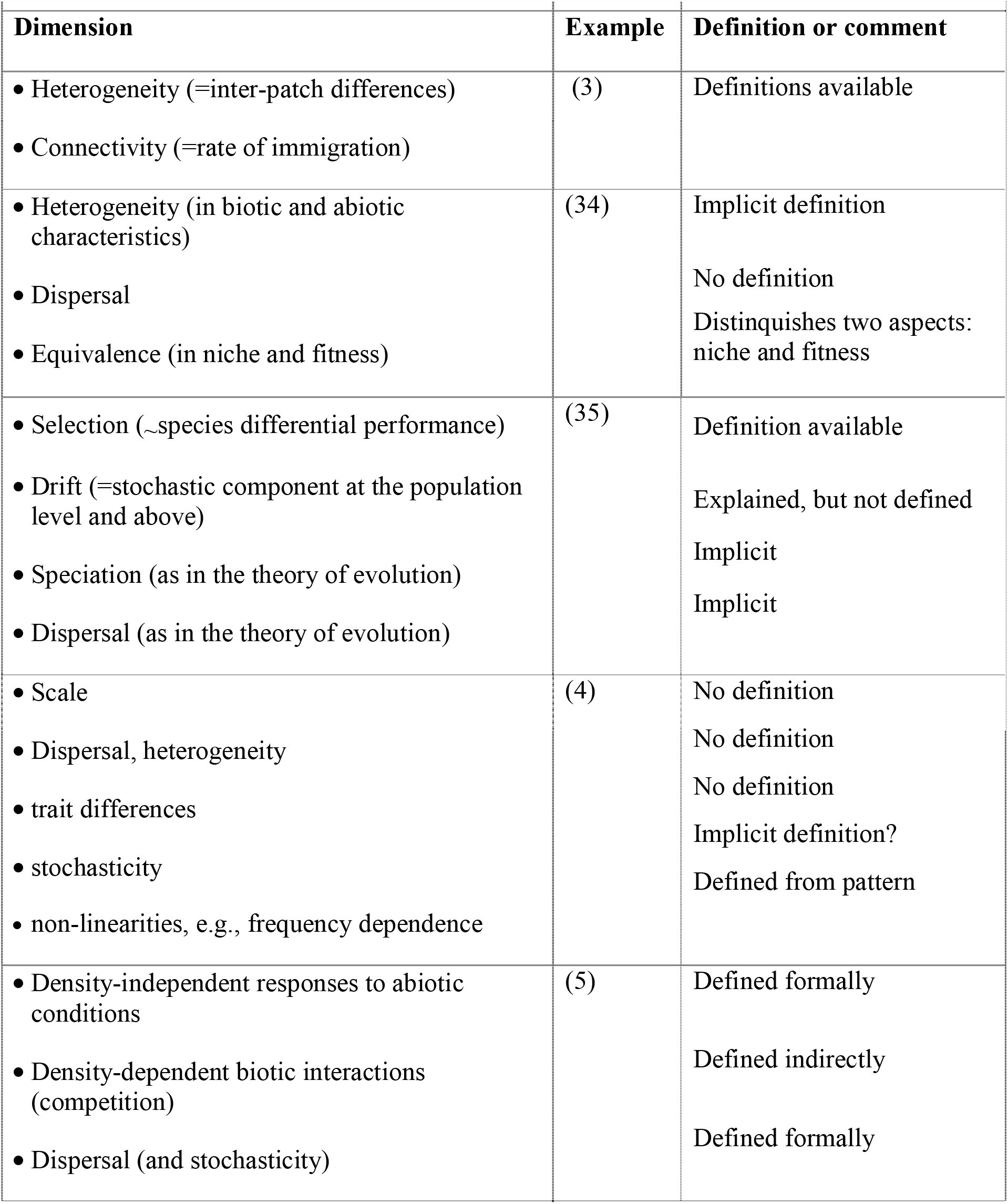
Examples of dimensions or related concepts for the MCF (only indirectly in Vellend, 2010). We note that some differences concern minor terminological conventions, whereas others involve swapping mechanisms with outcomes, studies on community dynamics usually ignored scales considerations, or introduce other unique approaches.

| Dimension | Example | Definition or comment |
| --- | --- | --- |
| • Heterogeneity (=inter-patch differences)<br>• Connectivity (=rate of immigration) | (3) | Definitions available |
| • Heterogeneity (in biotic and abiotic characteristics)<br>• Dispersal<br>• Equivalence (in niche and fitness) | (34) | Implicit definition<br>No definition<br>Distinguishes two aspects:<br>niche and fitness |
| • Selection (~species differential performance)<br>• Drift (=stochastic component at the population level and above)<br>• Speciation (as in the theory of evolution)<br>• Dispersal (as in the theory of evolution) | (35) | Definition available<br>Explained, but not defined<br>Implicit<br>Implicit |
| • Scale<br>• Dispersal, heterogeneity<br>• trait differences<br>• stochasticity<br>• non-linearities, e.g., frequency dependence | (4) | No definition<br>No definition<br>No definition<br>Implicit definition?<br>Defined from pattern |
| • Density-independent responses to abiotic conditions<br>• Density-dependent biotic interactions (competition)<br>• Dispersal (and stochasticity) | (5) | Defined formally<br>Defined indirectly<br>Defined formally |

**Table 2.**
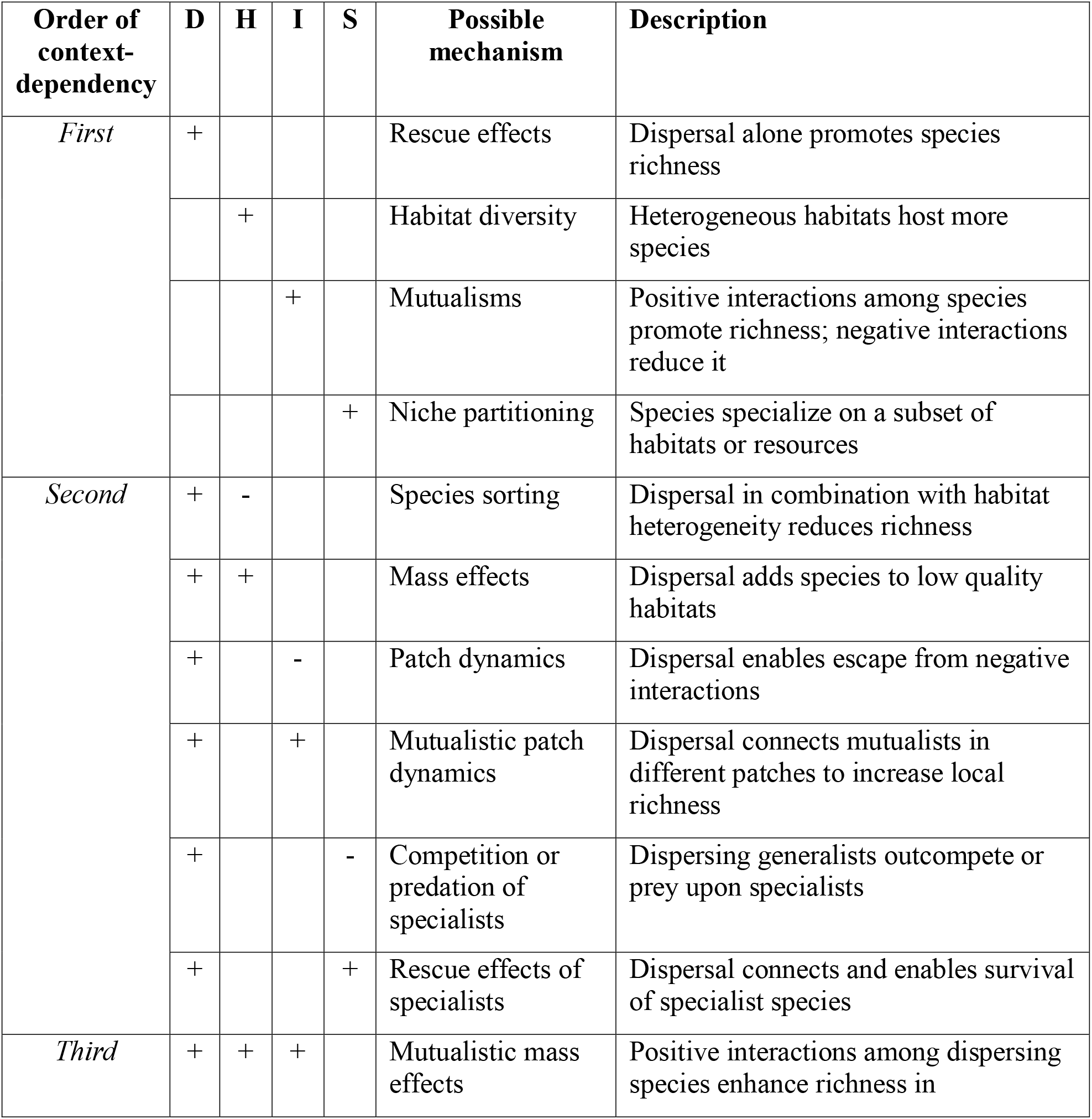
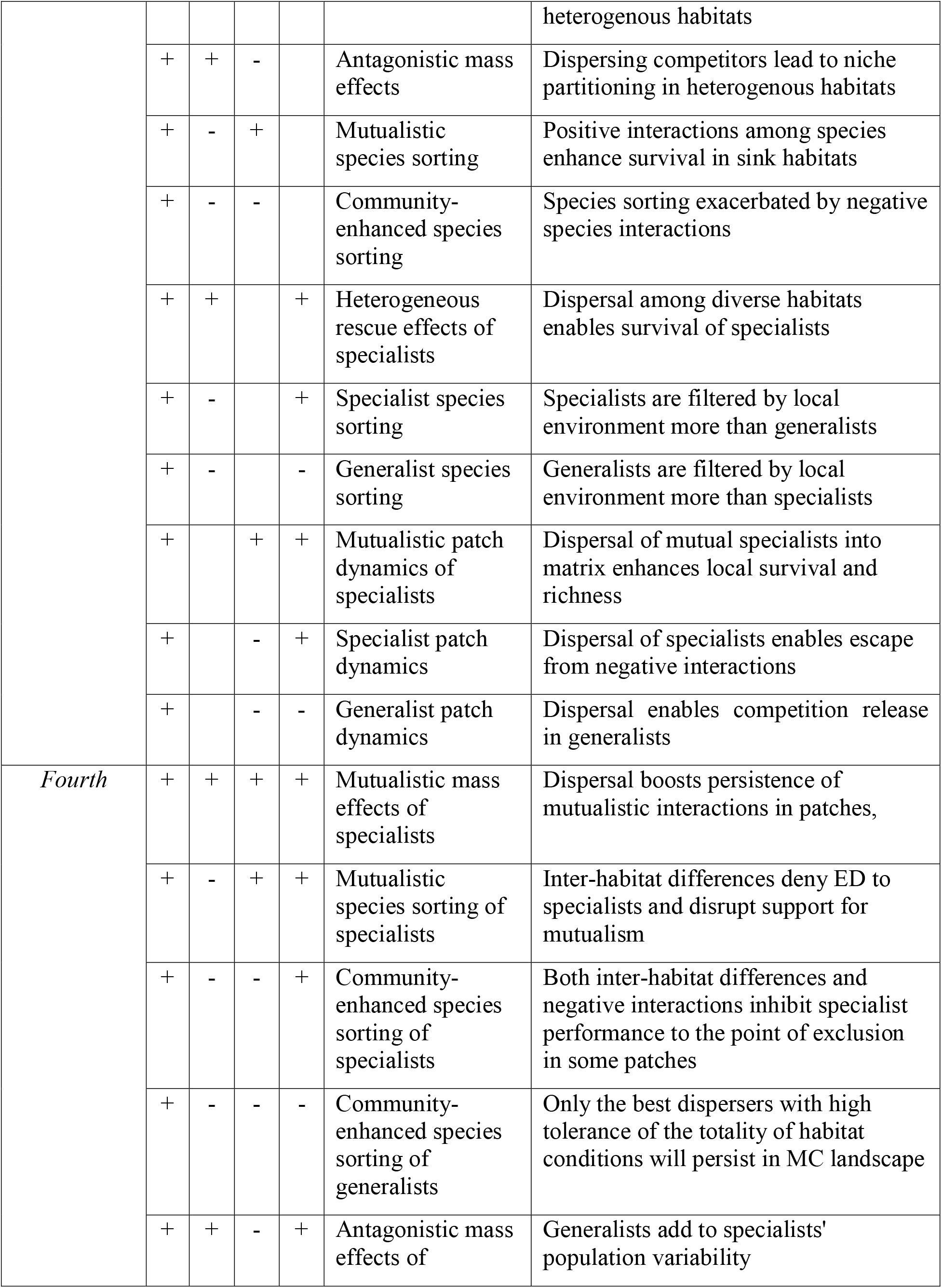
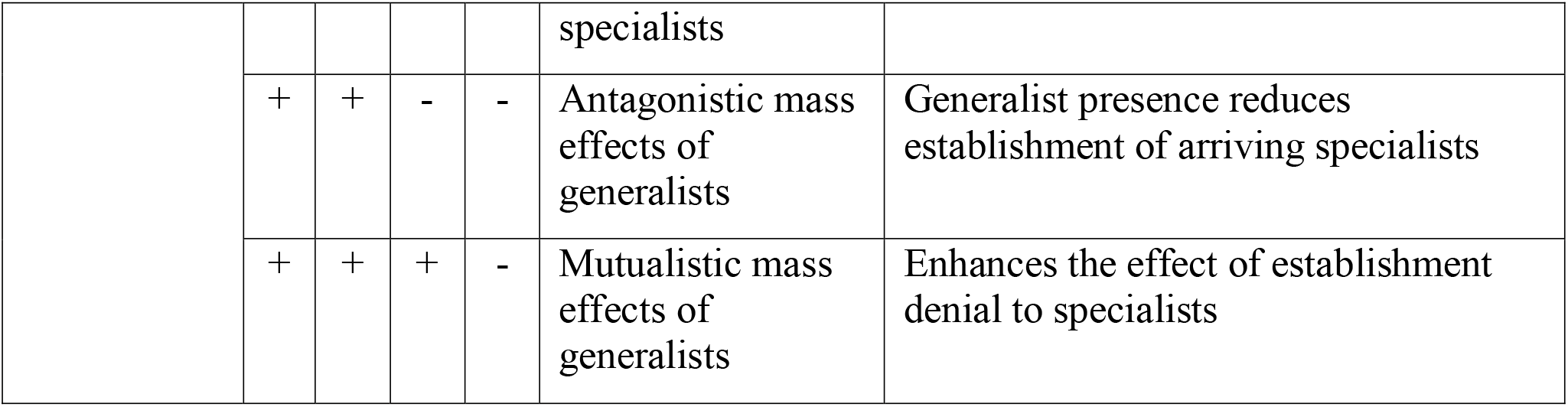
Interactions among dimensions – the raw material for hypotheses. 4D permutations underlying the context-dependency of metacommunity outcomes can serve as working hypotheses. D = dispersal; H = habitat heterogeneity; I = species interactions; S = specialization. + and − denote positive and negative effects of an individual factor on local species richness. The sign of factor interaction is given as the product of +s and −s. Only combinations including dispersal are presented to emphasize metacommunity processes.

Conceptual entanglement, CE, represents the second, less obvious, but likely more significant concern than terminological youth. Ecologists recognize that ecological phenomena frustrate the search for general laws and the formulation of pertinent theories when they are context-dependent or contingent ((9-11)). Familiarity with a standard answer to questions about outcomes: ‘it depends …’ is an epitome of context-dependency. While context-dependency is a consideration in most ecological studies, some forms pose only a modest challenge (Fig. 1, cases a-c) that empirical research tackles with appropriate experimental design, cf. (12), or statistics. One, however, affects the core of the theory.

**Fig. 1.**
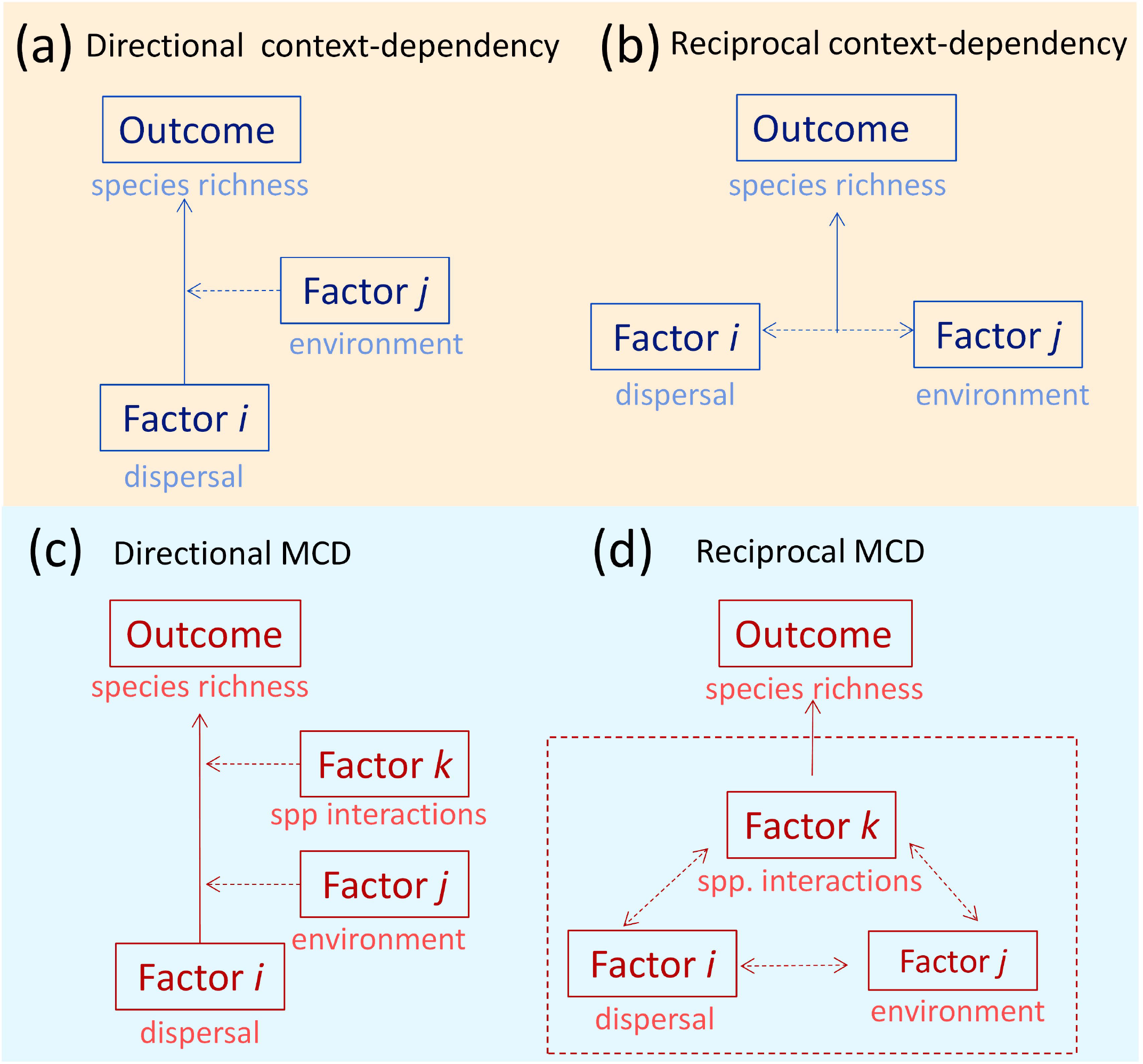
Basic ways in which multiple factors can combine into several types of context-dependency, CD (MCD – multiple CD), depending on how they interact. a-d differ in the number of factors and reciprocity of interactions (a and d – no reciprocity), b and c – reciprocity among two or more factors affects the outcome. Specific factors (dispersal, environment, interactions) appear as basic dimensions in MCF under similar labels.

When the interacting factors hold the status of the prime dimensions of an incipient theory, reciprocal dependency (MCD, Fig. 1d) represents a significant challenge. The predictions become ambiguous, entangled with three or more dimensions, and may fall anywhere within the inference space. This condition will likely decouple the theory from empirical tests because the tests rely on system-specific data (see (13, 14)). A failure to address this decoupling may render theory tests impractical or misleading unless a translation mode is available (10).

The many drivers of metacommunity dynamics may jointly determine an ecological dynamics and outcome (5), which poses conceptual and practical challenges (15). Context dependency continues to challenge our ability to predict the future states of communities and their ecosystem consequences (e.g., (15, 16)). The core problem is that where multiple variables jointly determine an outcome, an ecological process (e.g., dispersal) can result in appreciable differences when contexts differ. In short, inferences drawn in one setting do not apply in another unless we include other factors. This familiar adage of science requires considering multifaceted complexity (17). The geometric effects of many dimensions complicate likelihood of reaching reasonable predictions of system-level properties (14). Such systems allow more ways for pairwise outcomes to be different from ways to be similar. Despite the likelihood of considerable CD in metacommunities, theoretical treatments of CD are few and limited to single communities (e.g., (13). If it is also common in metacommunities, CD might thus impose severe limits on predicting biological outcomes from their different interacting processes unless we address it directly. Doing so requires a systematic approach.

### Approach: arranging MCF components

We propose that a hierarchical structure for MCF may mitigate the two problems, i.e., a) the proliferation of theory components and b) the entanglement of its dimensions. We chose four dimensions, 4D (Fig. 2), to provide the necessary and sufficient general structure for examining metacommunity processes and their outcomes.

**Fig. 2.**
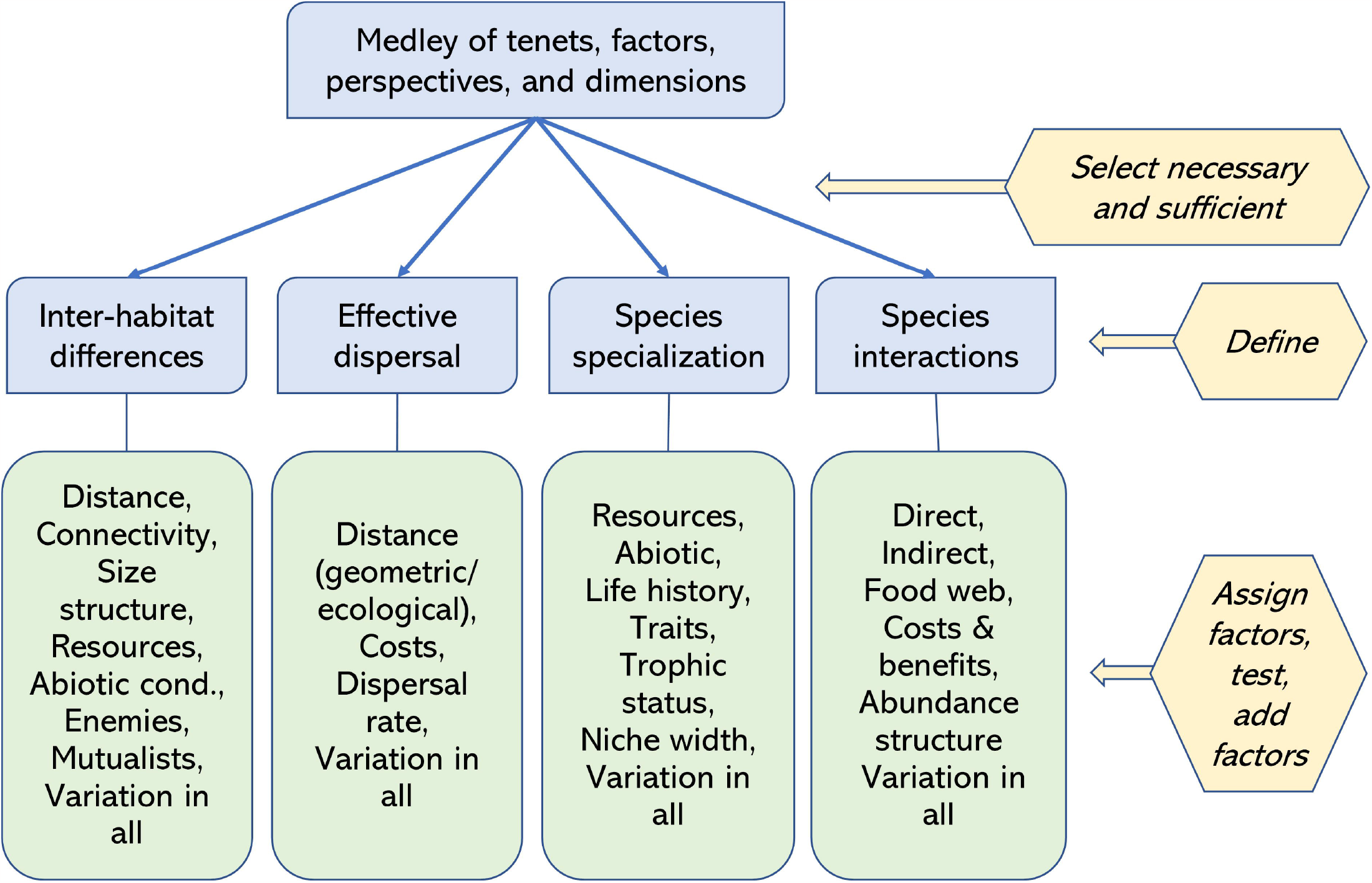
The general scheme. The top box acknowledges the diversity of propositions, models, approaches, and angles that appear in the literature (Table 1). Four dimensions can accomodate terms relevant to MCF. The scheme allows for adding other factors of interest. For example, position of a local habitat on a spatial gradient in a hierarchical system of river tributatries can appear as an inter-habitat difference or a modifier of the effective dispersal.

Species specialization combines species’ traits and habitat interactions. Species interactions include all possible effects species have on each other directly or mediated by habitat differences, dispersal, and species traits, cf. (18). Inter-habitat differences reflect species perceptions (cf. Fig. 3) for inter-dependencies among dimensions.

**Fig. 3.**
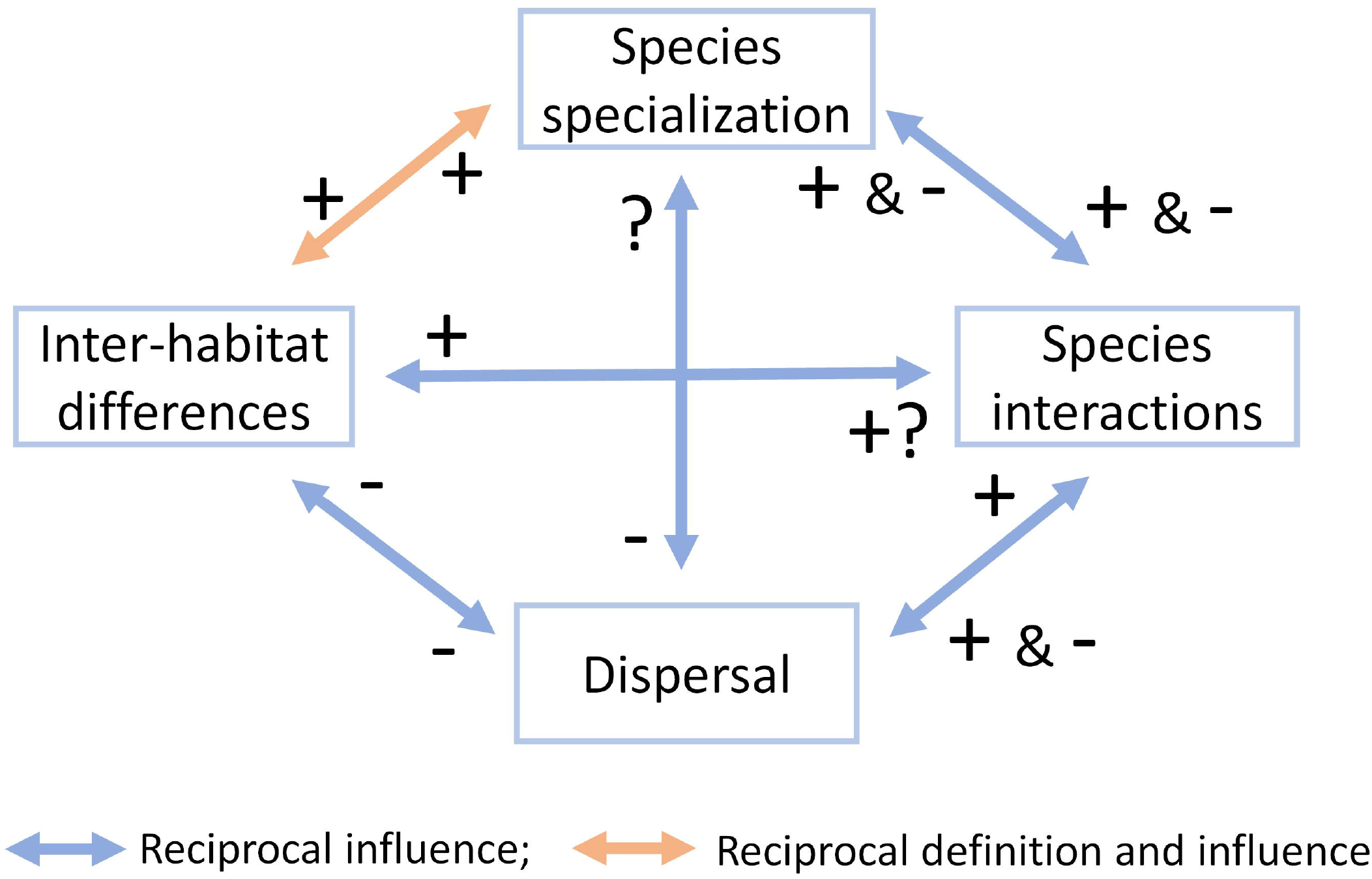
Known or expected reciprocal relationships among the four dimensions, 4D. Their effects may be positive or negative, direct and indirect. Further, we propose that species specialization and inter-habitat differences co-define each other (i.e., one cannot be meaningfully defined without the other) in addition to trading impacts. The signs indicate the most likely effect a value increase in one dimension has on another dimension.

We define the 4D, or macro-variables, in universal terms. The definitions allow for clear operationalization and can accommodate new variables as their sub-dimensions. Such growth mode allows for retaining the core MCF structure intact. We briefly explain 4D below:

a. **Species habitat specialization** (specialization hereafter) is the habitat space fraction a species can use. When no ecologically meaningful differences among patches exist (homogeneous habitat), species specialization remains undefined. In contrast, when inter-habitat differences are high enough to prevent species from crossing patch boundaries (maximum heterogeneity), all species become extreme specialists.
b. **Inter-habitat differences** are the second face of this relationship. The habitat is homogeneous when differences are insufficient to prevent species from differentially moving to and living in any patch. The Inter-habitat differences and Specialization show a positive reciprocal relationship: when one increases, the other increases.
c. **Species interactions** appear as a gradient from positive (+1) to negative (−1) without considering interaction types, cf. (19). Embedding them in the 4D system, underscores the vital modifying roles of other dimensions (other species interactions may also affect each specific species pair). Reciprocally, species interactions affect other dimensions by differentially changing the quality of habitat patches, redefining species specialization (restricting or expanding other species’ habitat access), and modifying effective dispersal.
d. **Effective dispersal** captures the dispersal process outcome (or a stage where its consequences begin). For example, an increase in effective dispersal diminishes interhabitat differences as perceived by individual species populations, with many secondary effects. We expect the effective dispersal to decline when interhabitat differences and specialization increase and thus to show a broader spectrum of effects in general (e.g., to increase local interaction intensity, modify their direction, or change food web connectivity (20)).

### Strategy: taking advantage of the many outcomes

The hierarchical structure of MCF (Fig. 2) and the inherent relationships among the high-level dimensions (Fig. 3) offer an opportunity ***to mitigate interference of context-dependency has on inferences***, at least when it comes to empirical testing of MCF predictions. First, we can determine the expected qualitative relationships among the 4D. This step will constrain the inference space and guide the prospective study design. Then, knowing the approximate position of the natural system on the gradient of possibilities, the relationship between observed and postulated patterns becomes easier to evaluate and compare across studies. Schematically, we can express the initial logic as:

> Perform coarse, high-level simulations → generate metrics analogous to those available in the study systems → identify the model configuration best matching observed metrics → formulate specific hypotheses → compare new output to observed metrics → and repeat the steps to uncover specific causal mechanisms until satisfied with the answer.

Below, we illustrate how a numerical model can reveal the potential of this strategy. We focus on patterns of species richness – a convenient metric of broad interest.

### Hypotheses

We center the further discussion on two particularly demanding CD types (Fig. 1c,d), with two corresponding hypotheses as tools for insights.

#### Type 1. Directional context-dependency

Here we test a simple prediction that ***holding a combination of parameters in three dimensions constant while changing values along one dimension will alter the effect of the remaining dimensions and thus the conclusion of a study (Hypothesis 1)***. We deduce this because the other dimensions can affect processes regulating local extinction and recolonization differently in response to the state of the focal dimension. Specifically, we envision that for any degree inter-habitat differences, the remaining dimensions may affect species richness enough to render habitat heterogeneity an inadequate predictor.

#### Type 2. Reciprocal context-dependency

A less common view of CD is that it emerges from interdependency among dimensions (e.g., (15). The reciprocal context-dependency arises when system dimensions entangle, i.e., when two or more factors affect each other non-additively in determining an outcome (Fig. 1b,d). Here, non-linear relationships and feedback among state variables may lead to an even richer network of interdependencies. Non-linearities may indeed be common in metacommunities (4). For instance, mixing distinct interspecific interactions can dramatically alter community dynamics and structure (e.g., (21). Thus, the dynamics, richness, and community composition predictions should be sensitive to slight changes in state variables or non-linearity in their relationships. Extreme sensitivity appears in simulated communities where the loss of patches needed for movement led to a cascade of diverse and unsuspected consequences mediated by a reduction in available habitat, connectivity, dispersal limitation, and other factors (22).

We assume that Reciprocal MCD (Fig. 1d) leads to a broader range of responses. We propose that: ***Context-dependency effects increase when fundamental dimensions of metacommunities can interact and influence one another* (Hypothesis 2)**. A test for this hypothesis may involve finding whether the effects of key variables are non-additive (change the sign or become non-linear) and lead to unexpected outcomes. Here, we examine how a confluence of antagonistic interactions, specialization, inter-habitat differences, and variability of specialist species impact regional species diversity. Revealing regularities in responses of dependent variables should allow for more accurate predictions and tests for specific systems.

### Specific features

We created the Unified Metacommunity Model, UMM, based on 4D, in the NetLogo software parameterized with the following features (details in Appendix 1):

**Habitat** - a mosaic of square patches in a regional meta-habitat with diverse suitability and resource (=energy) levels. Patch suitability values (fixed range of conditions that species must meet to use the patch) and resource quantities started as random. Resources regenerated at a random rate.

**Species** - individuals (agents) reproduce and move depending on their habitat requirements (randomly assigned) and the quality of the patch (suitability and state of resources) they occupy prior to the current step. The initial species richness was 50, the total abundance of 500 individuals, and the number of individuals a patch allowed was 25. A global carrying capacity of 920 individuals limited community population growth.

**Dispersal** - can occur when an individual acquires energy above a pre-set threshold. Movement direction was random, as was whether to disperse or reproduce. Individuals reaching a patch with available resources gained energy. The energetic cost of moving differed among species and species categories (e.g., specialists gain more from finding a suitable patch than generalists).

**Interactions** occur when an agent meets another agent(s) (including conspecific) on a patch. Interactions among species could be positive (mutualism, facilitation), negative (competition, predation, inhibition), or neutral (no direct impact or canceling indirect impacts). Interaction strength and outcome included parameters that gave specialists additional advantages upon meeting generalists or other specialists.

## Results

### Hypothesis 1

Empirical and theoretical work supports the idea that positive interactions improve environments for multiple species (19, 23) and can modify stability in model food webs [23]. Our simulations confirmed that specialization was a substantial factor in determining richness, but with a significant context-dependency: The richness-inflating effects of specialization hinged on positive interactions among specialists (Fig. 4A). The richness-promoting mechanisms we detect resemble those associated with species facilitation in general or symbiosis (24).

**Fig. 4.**
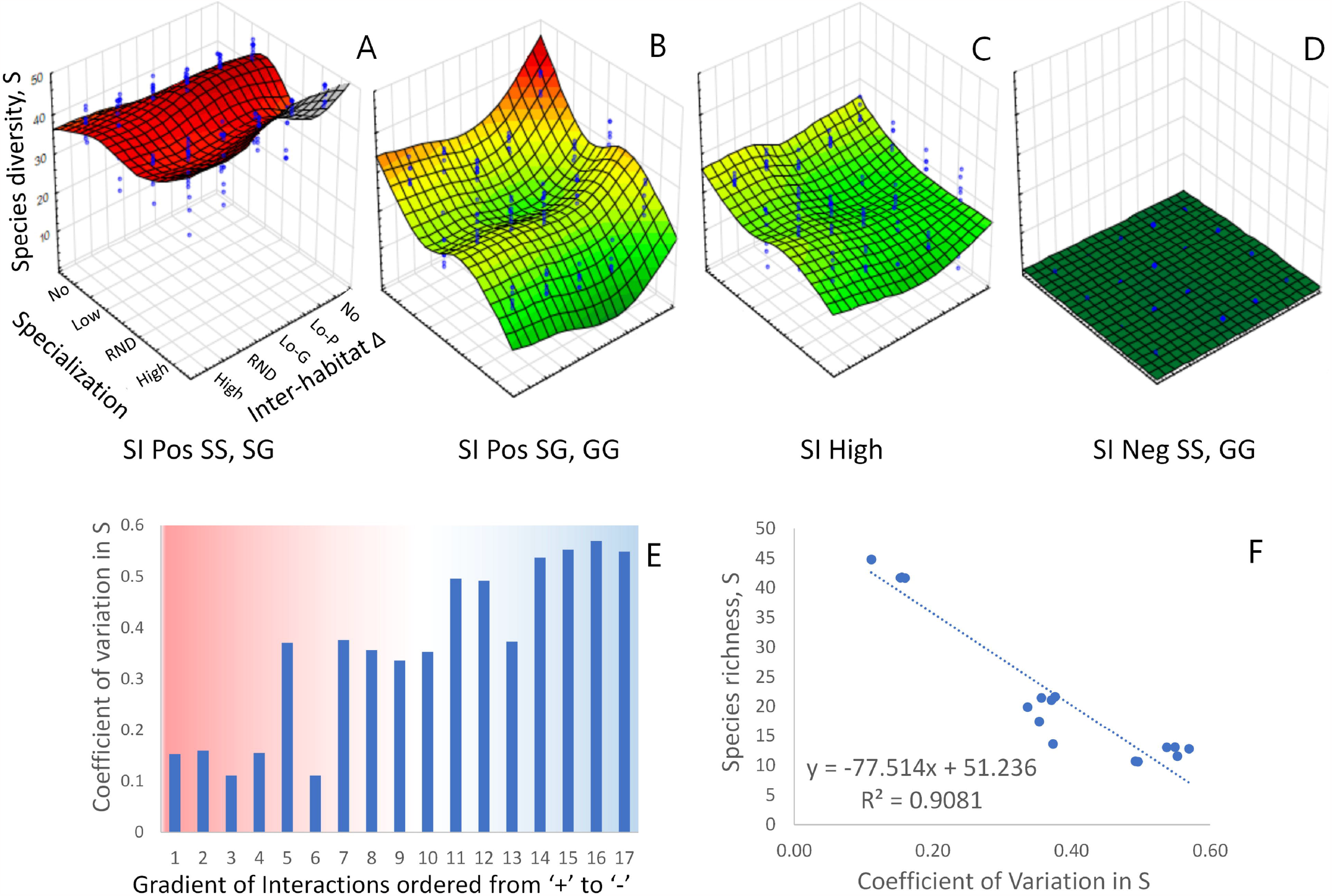
Different combination of interactions - on the gradient from negative to positive (a sample of all 18 available treatment combinations) - impact expected species richness, S (landscape mean). A-D – extreme positive (A) to negative (D) interaction configurations; B – the last positive, C – a random mix of positive and negative interactions; E – normalized variation increases with the shift towards negative interactions (from red to blue background) on a gradient of model treatments; F – the increase in variation is correlated with the loss of species richness. X-axes show Specialization ‘treatments’; y – axes show Inter-Habitat differences. Note differences in S and a loss of specialists near the middle range of species interactions. Pos – positive, Neg – negative, S – specialist, G-generalist, first letter – S or G affect the species S or G (second letter). Lo-G – low inter- habitat differences, good habitat; Lo-P - low inter-habitat differences, poor habitat. Complete set of results in Supplementary materials, Appendix 3, Fig. A3.1.

Interestingly, our simulations highlight the decisive role of positive interactions in boosting regional diversity when involving specialists. Specialists are typically rare and sensitive, e.g., (25), but contribute the most to species richness in a typical local community (26). Our observations converge with the idea that multiple ecological engineers (facilitators) increase stability by reducing primary extinctions and extinction cascade on community assembly in a model community (27). The previous study also found that species richness and stability increased with emerging interaction modules. In our simulations, higher variability in richness for negative over prevailing positive interactions (Fig. 4e) confirms that mutualism-like interactions boost metacommunity richness and stability, with habitat structure playing an indirect role.

Further, significant interactions among dimensions underscored that richness responses to one dimension were frequently modified by another (Appendix 2, for richness, S, Table A2.1 and abundance N_all_, Table A2.2). This pattern is diagnostic of the multivariate kind of CD in which multiple factors modify an ecological process (Fig 1c, d). Such reciprocal modifications imply that predictions based on incomplete sets of metacommunity dimensions, such as environmental gradients alone, will fall short of the mark. This also implies that conclusions from one context may fail to hold in another, even if locally correct. The problem may become acute when an influential process (e.g., specialized use of habitat) is assumed to have the same effect in another location or time, in circumstances where high context-dependency undermines the assumption, cf. (28).

### Hypothesis 2

Species richness of generalists and specialists responded differently to available resources, with resources becoming underutilized when specialist diversity declined (Fig. 5A). We also found that generalists and specialists richness responded differently to habitat suitability, with the latter having a hump-shaped response (Fig. 5B).

**Fig. 5.**
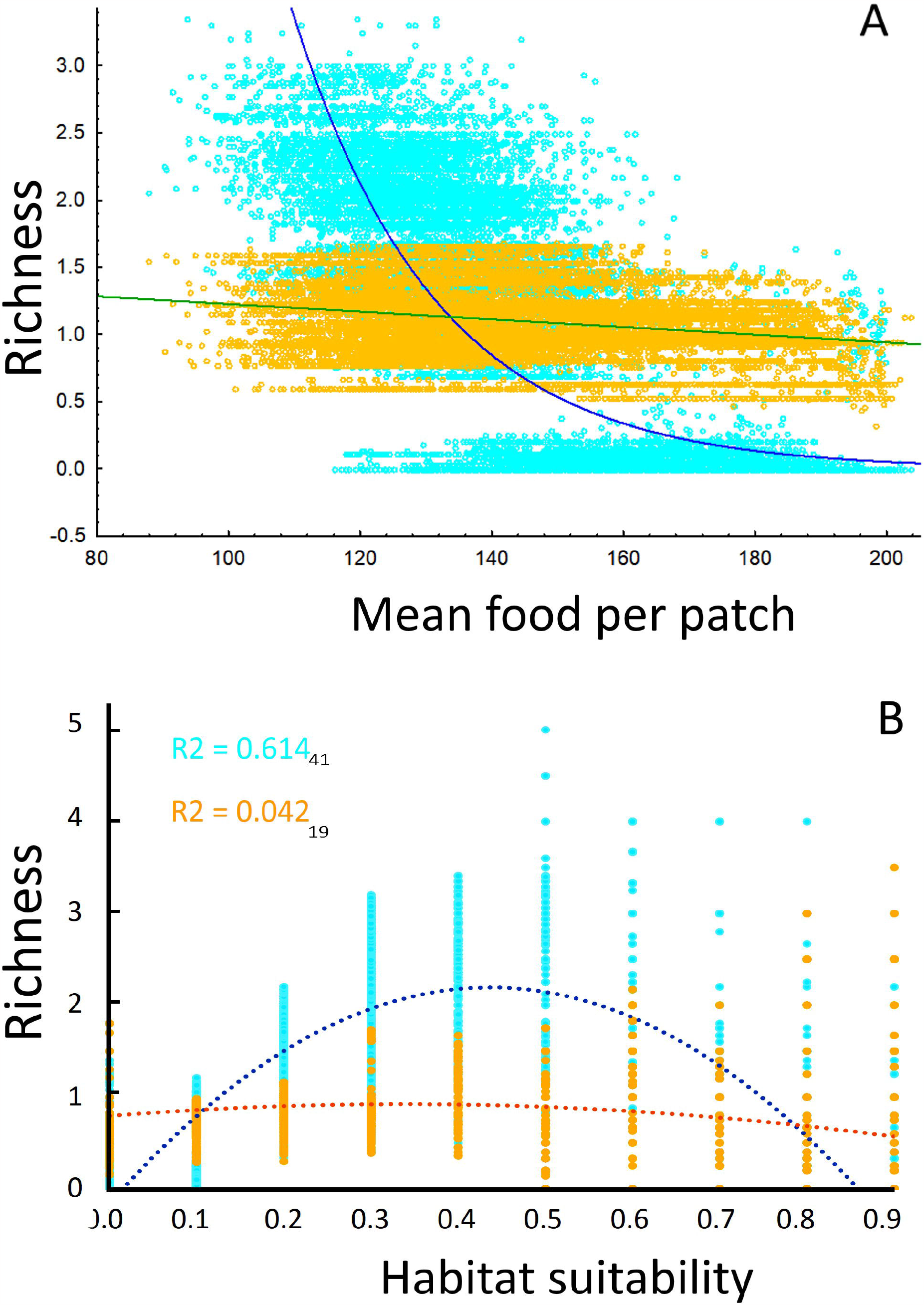
A - Species specialists (light blue) and generalist (goldenrod) richness, S, on the gradient of habitat suitability. B - Relationship between consumable resources and species richness of specialists and generalists. Diversity of specialists is associated with the amount of food resource (energy units in UMM), with a non-linear impact that is prominent in the middle of the habitat suitability range.

Indeed, neither food ‘causes’ declines in richness nor richness ‘causes’ declines in food. These two variables may correlate or not due to other processes. The UMM treats the entire food supply as available and consumed by all individuals present on a patch, irrespective of their species identity. Although patch energy regenerates in UMM, a patch does not compensate for energy consumption through recycling (e.g., (16). Other studies show that S may have complex relationships with density or biomass gradients (29).

To illustrate CD effects arising from the interplay between species interactions, species specialization, and habitat properties (inter-habitat differences), we chose treatment settings that yielded middle-of-the-range behavior of the UMM (similar to those in Fig 4C), with:

- High Specialization,
- Species Interactions where specialists benefitted from meeting other specialists and generalists while generalists suffered losses, and with
- Poor Habitat quality (low values of energy) and low Inter-Habitat differences.

We then tested whether the combined abundance of all species, N_all_, is a plausible explanation for the observed food decline. N_all_ related to Habitat Suitability in a curvilinear manner (Fig. 6A). This regularity was a mirror reflection of the amount of food available in each habitat suitability class (Fig. 5B). Relevant here is that the aggregated density of all species, N_all_, is negatively and linearly correlated with the amount of food on a patch (r^2^=0.917). In practical terms, species dispersal, successful recruitment, and habitat attractiveness changed directly or indirectly in response to each other. While revealing mechanisms contributing to this picture lies outside the scope of the paper, we note that species interactions appear to drive some of it. Specifically, we found that the combined population of specialists, N_spe_, initially increases, then decreases with N of generalists, N_gen_ (polynomial, quadratic regression: N_spe_ = −0.1877 N_gen_^2^ + 86.124 N_gen_ - 9519.5; R^2^ = 0.6154, n=40, p<0.00001). Thus, initial gains from the presence of generalists (cf. (27)) help boost specialists unless generalists increase in numbers and deplete resources – a pattern correlated with S_spe_.

**Fig. 6.**
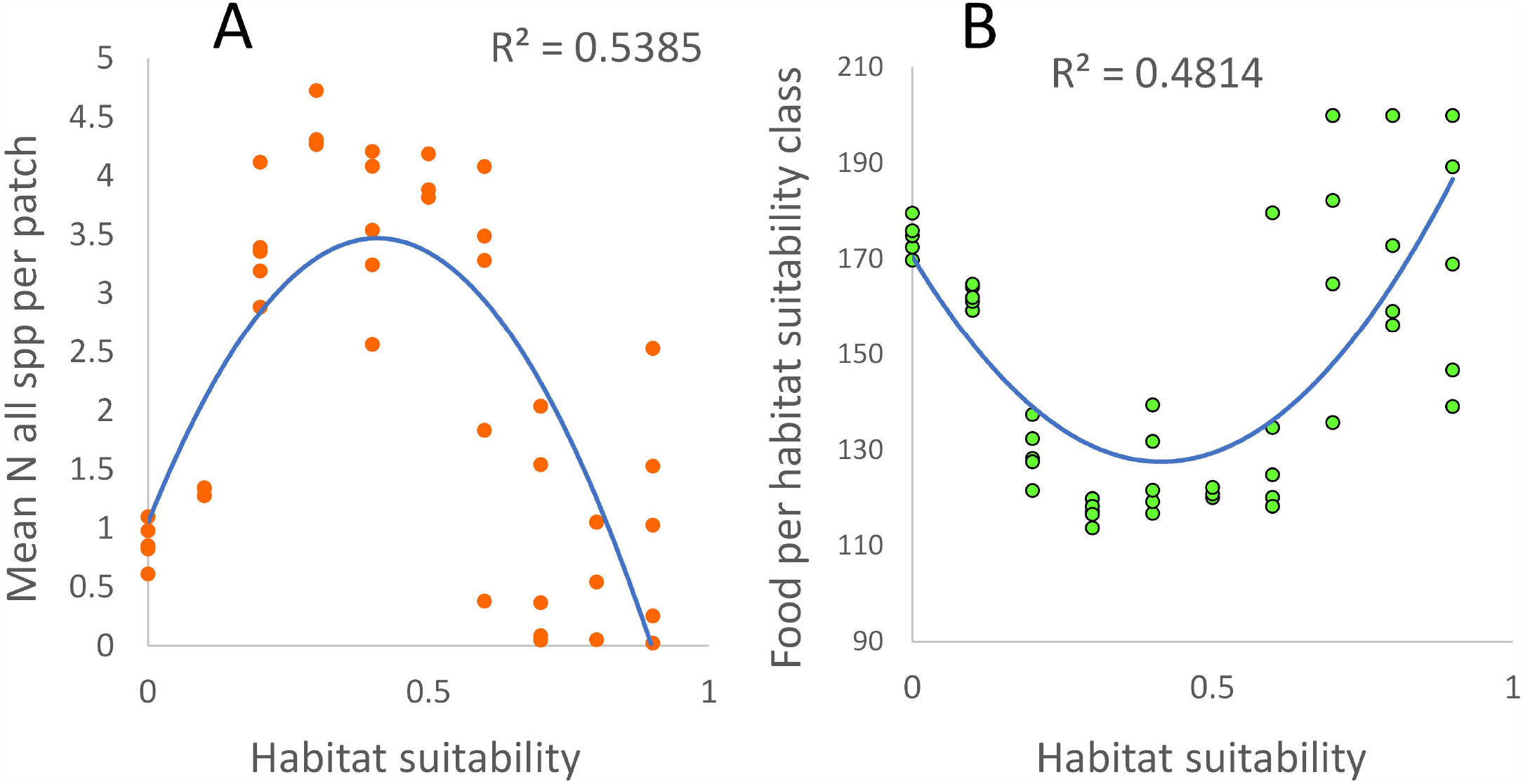
Mean aggregate abundance, N_all_, of species present (A) and food per patch suitability class (B) are related to habitat suitability class. Habitat suitability here represents initial ‘abiotic’ conditions that a species must satisfy. Total species abundance, N, correlates in a non-linear fashion with habitat suitability and provides functional explanation for specialist richness trend in Figure 4, but not for generalists.

## Discussion and conclusions

Our findings support that *reciprocal influences can affect other dimensions in the 4D framework and the metacommunity outcomes*. In our model exercise, a species’ success in reaching a patch, surviving, reproducing, and dispersing from it depends on two CD elements. One is compatibility with the non-interactive environment of the patch (abiotic and some biotic, e.g., trees and ground cover for ground rodents in a forest). Others are resources (represented as energy in the model) and species composition. These volatile aspects of heterogeneity can filter species distribution and local success. They can also trigger differential food consumption on patches with diverse suitability and accessibility values. In turn, this differential consumption modifies how species see/assess heterogeneity. These species-specific modifications translate into either dampening or amplifying inter-habitat suitability differences and modify the anticipated (e.g., in distribution, local and regional richness, population variation, and others, e.g., (30)). The classical study of intertidal invertebrates (31) highlights that environmental factors, species specialization, and interactions combine to produce spatial patterns difficult to predict from individual factors alone. The position of a habitat patch can also show significant CD in meta-habitats with a hierarchical structure, such as river networks (32, 33), and further modify environmental filters, dispersal, and species composition.

The practical result of reciprocal CD in metacommunities is that, even when accounting for essential dimensions, interdependencies, and reciprocities, feedback among them may lead to unexpected and emergent effects. Findings, therefore, challenge the tacit assumption that metacommunities are predictable from analyzing a few key dimensions. Because the net effects of interacting variables will be hard to predict at larger metacommunity or ecosystem scales, the prevalence of this type of CD will systematically undermine predictability. Even with knowledge of the system states in four dimensions, their interaction over time can give rise to ecological surprises.

The reasoning above may be familiar to succession or ecosystem researchers, but it is not integral to the metacommunity theory. MC research needs to espouse the view that the 4D dynamically adjusts and re-adjusts in response to each other in high-dimensional metacommunities. When such an adjustment occurs, it represents the most severe case of CD. In cases where individual dimensions form a dynamic network of interdependent processes, separating their contributions may be difficult to achieve in a single testing step, especially with incomplete data sets. We discuss separately some ways in which ecologists might recognize, mitigate, or even benefit from the entanglement of the MC framework constituent dimensions (Appendix 4).

## Acknowledgments

Natural Sciences and Engineering Research Council of Canada provided funding (Grant number 10531314), We are gratefulf to Drs. B. Brown, J. Orr, M. Leibold, R. Pringle, and S. Saavedra who provided insightful comments on the ideas and manuscript at various stages of the development, as did Drs B Fournier and Jinbao Liao, who reviewed the earlier version of the manuscript available on BioRxiv.

## References

1. Resetarits WJ, Binckley CA. Patch quality and context, but not patch number, drive multi-scale colonization dynamics in experimental aquatic landscapes. Oecologia. 2013;173(3):933–46.

2. Vepsalainen K, Spence JR. Generalization in ecology and evolutionary biology: From hypothesis to paradigm. Biology & Philosophy. 2000;15(2):211–38.

3. Leibold MA. The metacommunity concept and its theoretical underpinnings. In: Scheiner SM, editor. The theory of ecology. Chicago: The University of Chicago Press; 2011. p. 163–83.

4. Leibold MA, Chase JM. The Theories of Metacommunities. Metacommunity Ecology, Volume 59: Princeton University Press; 2018. p. 23–48.

5. Thompson PL, Guzman LM, De Meester L, Horvath Z, Ptacnik R, Vanschoenwinkel B, et al. A process-based metacommunity framework linking local and regional scale community ecology. Ecology Letters. 2020;23: 1314–29.

6. Winegardner AK, Jones BK, Ng ISY, Siqueira T, Cottenie K. The terminology of metacommunity ecology. Trends in Ecology & Evolution. 2012;27(5):253–4.

7. Brown BL, Sokol ER, Skelton J, Tornwall B. Making sense of metacommunities: dispelling the mythology of a metacommunity typology. Oecologia. 2017;183(3):643–52.

8. Guzman LM, Germain RM, Forbes C, Straus S, O’Connor MI, Gravel D, et al. Towards a multi-trophic extension of metacommunity ecology. Ecology Letters. 2019;22(1):19–33.

9. Guzman LM, Thompson PL, Viana DS, Vanschoenwinkel B, Horváth Z, Ptacnik R, et al. Accounting for temporal change in multiple biodiversity patterns improves the inference of metacommunity processes. Ecology. 2022;103(6):e3683.

10. Pickett STA, Kolasa J, Jones CG. Ecological understanding: the nature of theory and the theory of nature. San Diego, CA: Academic Press; 2007 2007.

11. Lawton JH. Are there general laws in ecology? Oikos. 1999;84(2):177–92.

12. Kimmel K, Dee LE, Avolio ML, Ferraro PJ. Causal assumptions and causal inference in ecological experiments. Trends EcolEvol. 2021;in press.

13. Song C, Von Ahn S, Rohr RP, Saavedra S. Towards a Probabilistic Understanding About the Context-Dependency of Species Interactions. Trends in Ecology & Evolution. 2020;35(5):384–96.

14. Orr JA, Piggott JJ, Jackson A, Arnoldi J-F. Why scaling up uncertain predictions to higher levels of organisation will underestimate change. bioRxiv. 2020:2020.05.26.117200.

15. Arnoldi J-F, Loreau M, Haegeman B. The inherent multidimensionality of temporal variability: how common and rare species shape stability patterns. Ecology Letters. 2019;22(10):1557–67.

16. Marleau JN, Guichard F. Meta-ecosystem processes alter ecosystem function and can promote herbivore-mediated coexistence. Ecology. 2019;100(6):1–11.

17. Fulton EA, Blanchard JL, Melbourne-Thomas J, Plagányi ÉE, Tulloch VJD. Where the Ecological Gaps Remain, a Modelers’ Perspective. Frontiers in Ecology and Evolution. 2019;7(424).

18. Poisot T, Canard E, Mouquet N, Hochberg ME. A comparative study of ecological specialization estimators. Methods Ecol Evol. 2012;3(3):537–44.

19. Koffel T, Daufresne T, Klausmeier CA. From competition to facilitation and mutualism: a general theory of the niche. Ecological Monographs. 2021;91(3):e01458.

20. LeCraw RM, Kratina P, Srivastava DS. Food web complexity and stability across habitat connectivity gradients. Oecologia. 2014;176(4):903–15.

21. Mougi A, Kondoh M. Diversity of Interaction Types and Ecological Community Stability. Science. 2012;337(6092):349–51.

22. Thompson PL, Rayfield B, Gonzalez A. Loss of habitat and connectivity erodes species diversity, ecosystem functioning, and stability in metacommunity networks. Ecography. 2017;40(1):98–108.

23. Stachowicz JJ. Mutualism, Facilitation, and the Structure of Ecological Communities: Positive interactions play a critical, but underappreciated, role in ecological communities by reducing physical or biotic stresses in existing habitats and by creating new habitats on which many species depend. BioScience. 2001;51(3):235–46.

24. Zélé F, Magalhães S, Kéfi S, Duncan A. Ecology and evolution of facilitation among symbionts. Nat Commun. 2018;9(4869):12.

25. Williams P. Does specialization explain rarity and decline among British bumblebees? A response to Goulson et al. Biological Conservation. 2005;122(1):33–43.

26. Kolasa J, Li B-L. Removing the confounding effect of habitat specialization reveals stabilizing contribution of diversity to species variability. Biology Letters. 2003;9:1–4.

27. Yeakel JD, Pires MM, de Aguiar MAM, O’Donnell JL, Guimarães PR, Gravel D, et al. Diverse interactions and ecosystem engineering can stabilize community assembly. Nat Commun. 2020;11(1):3307.

28. Kimbro DL, Byers JE, Grabowski JH, Hughes AR, Piehler MF. The biogeography of trophic cascades on US oyster reefs. Ecology Letters. 2014;17(7):845–54.

29. Duffy JE. Biodiversity and ecosystem function: the consumer connection. Oikos. 2002;99(2):201–19.

30. Petermann JS, Farjalla VF, Jocque M, Kratina P, MacDonald AAM, Marino NAC, et al. Dominant predators mediate the impact of habitat size on trophic structure in bromeliad invertebrate communities. Ecology. 2015;96(2):428–39.

31. Paine RT. Ecological determinism in the competition for space. Ecology. 1984;65:1339–48.

32. Mozzaquattro LB, Dala-Corte RB, Becker FG, Melo AS. Effects of spatial distance, physical barriers, and habitat on a stream fish metacommunity. Hydrobiologia. 2020;847(14):3039–54.

33. Dala-Corte RB, Becker FG, Melo AS. The importance of metacommunity processes for long-term turnover of riffle-dwelling fish assemblages depends on spatial position within a dendritic network. Canadian Journal of Fisheries and Aquatic Sciences. 2017;74(1):101–15.

34. Logue JB, Mouquet N, Peter H, Hillebrand H, Grp MW. Empirical approaches to metacommunities: a review and comparison with theory. Trends in Ecology & Evolution. 2011;26(9):482–91.

35. Kleynhans EJ, Otto SP, Reich PB, Vellend M. Adaptation to elevated CO2 in different biodiversity contexts. Nat Commun. 2016;7.

