## Supplementary information for "Metacommunity framework and its core terms entanglement"

### Appendices

Appendix 1

4D explained

The Unified Metacommunity Model (UMM) aims to expose how fundamental dimensions of metacommunities entangle to produce context-dependent outcomes (Fig. 2-3, main text).

The model's dimensions enable the testing of general postulates. However, as these dimensions are macro-variables, they also allow expansion to include subdimensions such as demographics, dispersal strategies, behavioral response, or configuration of patches. Therefore, the approach should encompass a range of reported mechanisms and dynamics.

We selected four dimensions, 4D, to organize processes of interest similar to those that already have a significant role in metacommunity research. They recognize that the movement of organisms in space, features of the space, traits of the organisms, and their interactions with each other are decisive determinants of various aspects of local and regional biodiversity. However, being macro-variables, the dimensions allow moving between general and specific questions and tests (Fig. 2, main text). They can either advance general postulates or explore specific aspects without missing the broader picture.

***Inter-habitat differences (heterogeneity)***

We define inter-habitat differences, IHΔ, as a function of species perceptions – here, a habitat is heterogeneous only to the extent to which differences among patches interfere with species growth rate, r_._ This dimension assumes continuous values on an arbitrary scale from 0 to 1, where 0 signifies a homogeneous matrix, and 1 signifies patch differences high enough to prevent species that occupies a single patch from occupying any other patch – a two-bookend-like case of counterfactual that defines the state space (34). In principle, IHΔ are based on the totality of conditions a patch presents to species of the regional pool and on barriers separating them, including distance. This definition of IHΔ implies that each species should experience landscape structure as different and that its experience changes with patch characteristics, including a complement of other species and available resources on the patch. For instance, species distributions in Gorongosa National Park (Box 2) relate to organism perceptions of how dangerous diverse habitat types are. Removal of predators reduced heterogeneity in those perceptions as predators define some areas as more dangerous than others (35, 36). Importantly, this change occurs even if habitat heterogeneity perceived by an observer has not changed.

***Dispersal***

We base *effective dispersal* rate, or just dispersal rate (37, 38), on the *probability of colonizing* rather than general mobility – the number of individuals leaving a source patch individuals dispersing, or moving (sensu (8). We chose this metric because it directly focuses on individuals added to and participating in a local network of community processes. Mobility matters but may depend on short-term patch conditions (positive and negative interactions, resources, abiotic quality).

As species activities modify these conditions, mobility may elevate the uncertainty of predictions. The interpretation of spatial transfer of individuals that we adopt here ties the effective dispersal to IHΔ above and, indirectly, to species composition in patches that species use as steppingstones to the final receiving patch. For example, suppose positive interactions dominate a patch, and the mean specialization of community members is high. In that case, effective dispersal into that patch will increase S until the community reaches its carrying capacity while remaining undefined prior to it. This view opens a new avenue for questions and hypotheses.

***Specialization***

We treat specialization as a breadth of species response to a collection of diverse habitats. This view combines the effects of habitat properties, species traits, predators, competitors, mutualists, and other factors. In the model, a species' distribution on a gradient of inter-habitat differences serves as a metric of specialization. Because species response is context-dependent and often non-linear (39), we propose a modified approach to account for the role of traits in MC dynamics. The approach partially overlaps with the popular version of the realized niche concept, including the niche-habitat relationship developed by (40). The meaning of specialization we propose here differs from the niche-based formulations.

Here, the specialization is contingent on the habitat structure (inherent adaptability), making it scalable across varying levels of heterogeneity. Further, it reflects history and priority effects and combines the effects of abiotic (including space) and biotic interactions (including indirect ones (cf. (5)). In general, it helps with two issues.

- It helps attain a concise list of general dimensions.
- It blends the distinction between biotic density-dependence and responses to abiotic factors. This distinction is not as obvious or necessary as standard references might suggest. Notably, when these two categories of factors interact are often used as one (41).

***Species Interactions***

A general view of MCF requires accommodating its most recognized feature - competition cf. (5, 32, 37), other direct interactions such as predation (42-47), mutualism (48), parasitism, commensalism, as well as a high number of possible indirect interactions (e.g., (49-51). Although most studies focused on a single or few interaction types, an interaction network active in a natural MC is likely to involve many. Extending the MCF approach to ecosystem properties makes this network even more complex (e.g., (52) and produces correspondingly revealing and unexpected outcomes such as emergent patterns of interactions, non-linearities (53), and spontaneous self-organization (54, 55), along with habitat modifications.

To accommodate a wide range of ecological interactions at the highest level of MCF, we define interactions as a generalized impact of one species on another. This impact is scaled from -1 to 1, with values around 0 corresponding to no interaction, to accommodate commonly discussed direct (competition, predation, mutualism, parasitism) and indirect interactions, including those mediated by the environment (e.g., creosote bush that facilitates the growth of saguaro cactus via microclimate amelioration).

The UMM design allows interactions among the four core metacommunity dimensions, represented by the arrows in Figure 3. These links reveal some already recognized forms of CD. Possibly the earliest recognized of these is dispersal-induced interactions where the sign or magnitude of species interactions depends on dispersal, such as in prey escaping predation.

Another known form of CD is filtered interactions, where the local environment filters the community to a subset of species and modulates their interactions (56); (57). Other forms are less recognized. Condition-dependent dispersal is when its rate depends on species interactions, such as strong local competition (58). The contingencies possible in MCs go beyond pairwise interactions and involve three or four dimensions (Table 2).

### Appendix 2: Methods

#### Unified metacommunity model, UMM

We implemented a simulation model within NetLogo agent-based framework, ver 6.1.1, which allows defining explicit patchy environments and species with different traits living and interacting with each other and their environment. To best answer questions about CD, the model is process-based, with elements of stochasticity to at the level of individual agent (an individual of a species) behavior, specialization, and abundance. Initial patch quality and patch arrangement are also stochastic within predetermined bounds.

**Structure - species**

Each of the four dimensions discussed in the paper represents a range of categories that span a spectrum of realistic situations seen in natural systems. We used 50 species with randomly generated traits either across the complete set of species or within categories of specialization and inter-specific interactions. In each replicated run, UMM assigns species identities and traits anew.

**Structure - Environment**

A no-edge landscape mosaic of 81 patches represents the environment. The position of patches of different quality was random for each replicated run. Heterogeneity implementation involved four selectable options (see Inter-habitat differences below).

**Individual Dimensions**

###### ***Specialization*** included *Low, High, Random, and None*. Species are assigned different ranges of specialization that determine which patches they can use. A broader range of specialization gives a species access to more patches, which reduces dispersal costs and increases the amount of resources that species can access. For example, the setting *High* creates three contiguous (with one minor discontinuity) classes of species, two of which have narrow specializations (vary from >0 to 0.2 and from >0.2 to 0.4, respectively) and a class of generalists (from >0.5 to 1). The latter class comprises 10% of species only.

***Inter-habitat differences***, IHΔ, (heterogeneity) included *High, Low-Good, Low-Poor, and None*. These differences were a function of randomly assigned *Suitability* values in a range of 0, 1 (mean 0.5) with the following modifications. For the *High* option, a patch was allowed a random value from the entire range (mean 0.5); for *Low*, the range was restricted to 0, 0.5 (mean of 0.325); for *Low-Good,* the range was 0.6, 1 (mean 0.8).

***Dispersal*** included: *High, Non-Limited, Random, Low*, and *None* rates defined by a fraction of agents (individuals) randomly chosen from all individuals present in the landscape).

***Interspecific interactions*** among species) included positive, negative, and neutral. Positive interaction gives an individual of a species an energy supplement, negative results in energy loss, and neutral does not cause an energy change. Interspecific interaction allows for indirect effects mediated by the energy supply a patch had (diffused competition) and by presence of intermediate interactors (a positive effect of one species on another may result in a negative effect on individuals of the third species when the initial beneficiary reproduced using energy from the first interaction).

**The Code**

The model code v.13 is/will be publicly available from NetLogo Community Models: [NetLogo User Community Models (northwestern.edu)](https://ccl.northwestern.edu/netlogo/models/community/). The model explanation (PowerPoint) is available here (when the manuscript is accepted).

#### Data and analyses

We ran UMM choosing one set of factors from 600 combinations allowed under the 4D and their categories. Each combination was replicated 10 times and the means recorded. We then assembled those means in a single data file. The data file was analyses with multi-factorial module, Statistica v. 13.5.0.17.

Additional simulations to examine links between richness, S, of specialists and generalists and the habitat resources (Supplementary materials, Results, Fig. A3.1) involved 250 runs in five replicates of the following settings (Supplementary materials, Appendix 1, Table A2.1):

***Table A2.1****. Explanation of settings. Species interactions included energetic benefits accorded to Specialists in gs-, gg-, ss- and sg-reward schedule (0 or negative value for generalists, +10 or 0 value for specialists). Inter-habitat differences were Low and the Habitat suitability was Poor on average. Species specialization was high (only few species allowed to use rand of habitats spanning >0.5 and the remaining species were assigned habitat range use <0.5). Effective dispersal High means that all individuals of a species can move 2 habitat units. Patch Regrowth rate allows food resource on a patch to drop to 50% of the starting value and then replenishes it at a set rate.*

| Species interactions High | Reproduction-rate 0.8 |
| --- | --- |
| Interhabitat differences Low-poor | Boost-reproduction? False |
| Species specialization High | Initial-individuals 500 |
| Effective dispersal High | Initial-food-per-patch 200 |
| Number-of-species 50 | %-generalists N 0.36 |
| K-always-on? True | K fraction 0.75 |
| gs-reward 10 | Specialists S 47 |
| gg-reward -10 | ss-reward 0 |
| Patch-regrowth-rate 0.96 | sg-reward 0 |
| Num-runs 250 |  |

### Appendix 3: Graphical and analytical patterns

### UMM dimensions have major impacts on S (Table A3.1), with abundance as a continuous covariate. Dispersal does affect S only, sometimes marginally, when in combination with normalized variation of specialists as continuous covariate (Table A3.2). This effect seems to be related to how habitat mosaic impedes specialists' movements and enhances stochastic variation of species, with colonization options depending on a current configuration of good/poor suitability patches.

**Table A3.1**. An example of variability in model outcomes where species richness S depends on the choice of interspecific interaction categories. Case 1: All 4 D used (see Methods above) with Species Interactions selected to be Specialists-Generalists (many positive interactions favoring specialist species). Case 2: Variable selection is the same as in Case 1, except that Species Interactions are random. Case 3: Species interactions are dropped entirely while the remaining variables are the same as in two previous cases. General linear model - Factorial ANOVA, n = 3600. Abbreviations: IH**Δ** – Inter-habitat differences; SI – Species interactions; Significant terms are bold (text) or red (numbers). Note difference in the number and type of significant terms among cases. Case 3 suggests that deducing factors promoting species richness when species interactions are unknown would be misleading. Equivalent results were observed for other individual dimensions. Red font highlights significant relationships.

| **Effect** | **SS** | **DF** | **MS** | **F** | **p** |
| --- | --- | --- | --- | --- | --- |
| 1. **All 4D, Species interactions = Specialists and Generalists** |  |  |  |  |  |
| **Intercept** | 1794350 | 1 | 1794350 | 61421.11 | 0.0000 |
| **Specialization** | 3470 | 3 | 1157 | 39.59 | 0.0000 |
| **IHΔ** | 7066 | 4 | 1766 | 60.47 | 0.0000 |
| **Dispersal** | 1626 | 4 | 406 | 13.91 | 0.0000 |
| **SI (Spec-Gen)** | 678829 | 17 | 39931 | 1366.85 | 0.0000 |
| **Specialization*IHΔ,** | 1180 | 12 | 98 | 3.37 | 0.0001 |
| Specialization*Dispersal | 215 | 12 | 18 | 0.61 | 0.8325 |
| **IHΔ*Dispersal** | 1169 | 16 | 73 | 2.5 | 0.0009 |
| **Specialization*SI (Spec-Gen)** | 47978 | 51 | 941 | 32.2 | 0.0000 |
| **IHΔ, *SI (Spec-Gen)** | 12466 | 68 | 183 | 6.27 | 0.0000 |
| **Dispersal*SI (Spec-Gen)** | 4162 | 68 | 61 | 2.1 | 0.0000 |
| Specialization*IHΔ, *Dispersal | 251 | 48 | 5 | 0.18 | 1.0 |
| Specialization*IHΔ, *SI (Spec-Gen) | 2984 | 204 | 15 | 0.5 | 1.0 |
| Specialization*Dispersal*SI (Spec-Gen) | 1416 | 204 | 7 | 0.24 | 1.0 |
| IHΔ, *Dispersal*SI (Spec-Gen) | 2532 | 272 | 9 | 0.32 | 1.0 |
| Specialization*IHΔ, *Dispersal*SI (Spec-Gen) | 4539 | 816 | 6 | 0.19 | 1.0 |
| 2. **All 4D, Species interactions = Random** |  |  |  |  |  |
| **Intercept** | 351174.0 | 1 | 351174.0 | 1455.10 | 0.0000 |
| **Specialization** | 3140.7 | 3 | 1046.9 | 4.34 | 0.0047 |
| IHΔ | 1822.4 | 4 | 455.6 | 1.89 | 0.1098 |
| Dispersal | 701.2 | 4 | 175.3 | 0.73 | 0.5738 |
| **Interactions (RND)** | 6510.5 | 2 | 3255.2 | 13.49 | 0.0000 |
| Specialization*Interactions (RND) | 1147.3 | 6 | 191.2 | 0.79 | 0.5758 |
| IHDelta, *Interactions (RND) | 1079.0 | 8 | 134.9 | 0.56 | 0.8122 |
| Dispersal*Interactions (RND) | 496.7 | 8 | 62.1 | 0.26 | 0.9791 |
| Specialization*IHΔ, *Dispersal | 384.5 | 48 | 8.0 | 0.03 | 1.0 |
| Specialization*IHΔ, *Interactions (RND) | 201.3 | 24 | 8.4 | 0.04 | 1.0 |
| Specialization*Dispersal*Interactions (RND) | 377.8 | 24 | 15.7 | 0.07 | 1.0 |
| IHDelta, *Dispersal*Interactions (RND) | 390.4 | 32 | 12.2 | 0.05 | 1.0 |
| Specialization*IHΔ, *Dispersal*Interactions (RND) | 866.7 | 96 | 9.0 | 0.04 | 1.0 |
| 3. **3D – no Species interactions** |  |  |  |  |  |
| **Intercept** | 1794350 | 1 | 1794350 | 7777.463 | 0.0000 |
| **Specialization** | 3470 | 3 | 1157 | 5.013 | 0.0018 |
| **IHΔ** | 7066 | 4 | 1766 | 7.657 | 0.0000 |
| Dispersal | 1626 | 4 | 406 | 1.762 | 0.1337 |
| Specialization*IHΔ | 1180 | 12 | 98 | 0.426 | 0.9539 |
| Specialization*Dispersal | 215 | 12 | 18 | 0.078 | 1.0 |
| IHΔ *Dispersal | 1169 | 16 | 73 | 0.317 | 0.9953 |
| Specialization*IHΔ *Dispersal | 251 | 48 | 5 | 0.023 | 1.0 |

**Table A3.2**. Univariate Tests of Significance for Total N (individuals). The 4D are cell-centered, bold type indicates significant relationships, p<0.000, Sigma-restricted parameterization, effective hypothesis decomposition; Std. Error of Estimate: 78.45. The red font highlights significant relationships.

| **Effect** | **SS** | **DF** | **MS** | **F** | | | **p** |
| --- | --- | --- | --- | --- | --- | --- | --- |
| **Intercept** | 6.74E+08 | 1 | 6.74E+08 | | 72659.81 | 0.0000 | |
| **Specialization** | 2737593 | 3 | 912531 | | 98.42 | 0.0000 | |
| **Interhabitat Δ** | 1005566 | 4 | 251391 | | 27.11 | 0.0000 | |
| **Dispersal** | 174275 | 4 | 43569 | | 4.7 | 0.0019 | |
| **SI(Spec-Gen)** | 1.19E+08 | 17 | 7011280 | | 756.23 | 0.0000 | |
| Specialization*Interhabitat Δ | 114751 | 12 | 9563 | | 1.03 | 0.4165 | |
| Specialization*Dispersal | 79107 | 12 | 6592 | | 0.71 | 0.7420 | |
| Inter-habitat Δ *Dispersal | 37603 | 16 | 2350 | | 0.25 | 0.9988 | |
| **Specialization*SI(Spec-Gen)** | 1083353 | 51 | 21242 | | 2.29 | 0.0000 | |
| Interhabitat Δ*SI(Spec-Gen) | 774639 | 68 | 11392 | | 1.23 | 0.1020 | |
| **Dispersal*SI(Spec-Gen)** | 2446098 | 68 | 35972 | | 3.88 | 0.0000 | |
| Specialization*Interhabitat Δ*Dispersal | 19703 | 48 | 410 | | 0.04 | 1 | |
| Specialization*Interhabitat Δ*SI(Spec-Gen) | 109115 | 204 | 535 | | 0.06 | 1 | |
| Specialization*Dispersal*SI(Spec-Gen) | 135929 | 204 | 666 | | 0.07 | 1 | |
| Interhabitat Δ*Dispersal*SI(Spec-Gen) | 186958 | 272 | 687 | | 0.07 | 1 | |
| 1*2*3*4 (all 4D) | 301625 | 816 | 370 | | 0.04 | 1 | |
| Error | 16688467 | 1800 | 9271 | |  |  | |

Eithghteen combinations of species interactions with the three remaining dimensions, 3D, show variation in outcomes. This variation forms a broad gradient with some degree of similarity among patterns emerging in adjacent areas of inference space (Fig. A3.1).

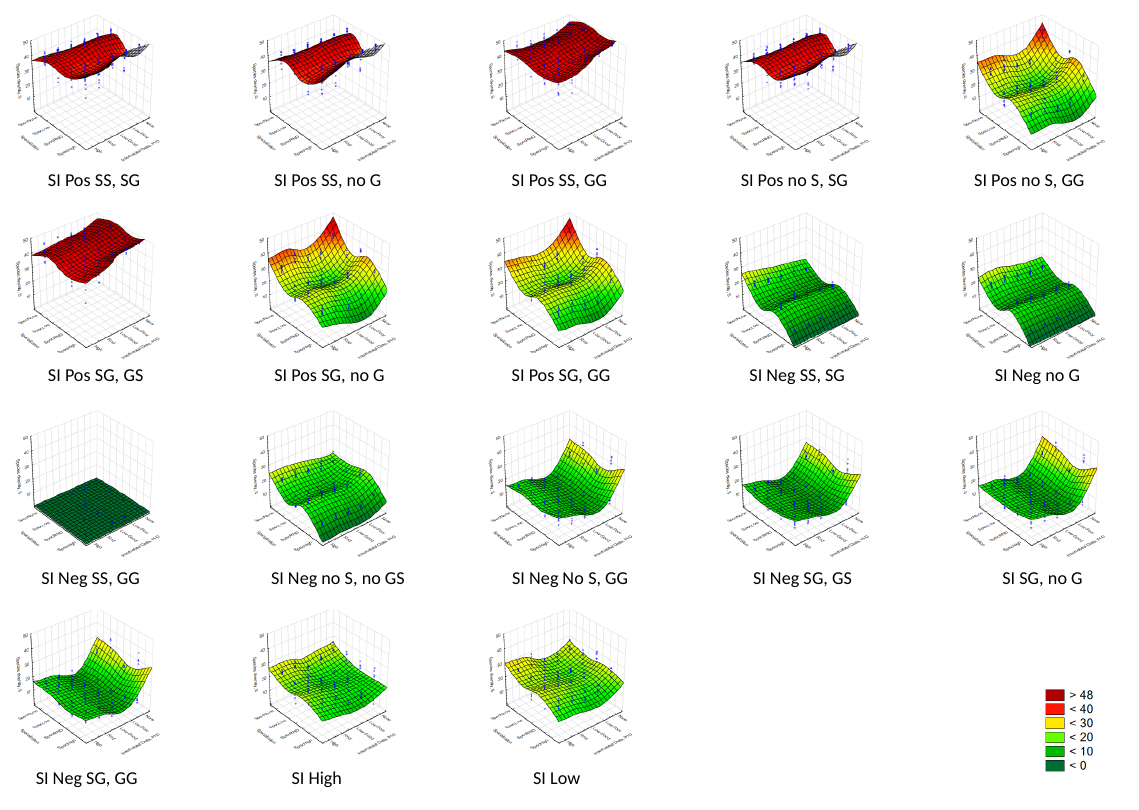

*Figure A3.1. Different combinations of species interactions on the gradient from positive to negative affect a metacommunity metric: species richness, S (landscape mean). The first letter – S or G – identifies the direction of the effect from the donor on the receiver species, S or G (second letter). Y-axis – species diversity, S; x-axis (left) – species specialization; and z-axis – inter-habitat delta categories (all readable upon a zoom-in). Axes labels as in the main text Figure 4A.*

### Appendix 4: Dimension entanglement in the future metacommunity research

##### Implications. Recognizing the entanglement of the fundamental dimensions is the first step on the path to a resilient and general theory. The example of MC dynamics along one dimension revealed a substantial shift among patterns (Fig. 4). These patterns may be sensitive to small changes along other dimensions that are not included in analyses, especially where species manage to modify habitat quality (Fig. 6). While researchers understand the challenge of predicting outcomes in species-rich landscapes, CD is yet to become an integral part of metacommunity theory and should be easier after the entanglement among 4D are fully understood. When we allowed species to modify habitat quality, prominent differences among species of different specializations (Fig. 4) emerged, along with distinct non-linearities in other pivotal variables (Fig. 5). Together, they warn against a simple research strategy. Nevertheless, we see a realm of general possibilities around which a more calibrated strategy might grow (cf. Fig. 2, main text):

1. In low-dimensional systems, one dimension may suffice for prediction.

2. In high-dimensional (or complex) systems:

a. Linear models of the 4Ds may detect directional CD (cf. Fig. 1c), i.e., models capable of capturing the main and modifying effects of separate dimensions.

b. Detecting and correcting for the reciprocal CD (cf. Fig. 1d) will be challenging to detect because of non-linearities arising from "back-and-forth" among the dimensions. To quickly explore generic behaviors as opposed to a strategy of testing a limited number of models against convenient natural systems, dynamic models may help.

The above suggests that remedies will require recognizing where CD plays a significant role and designing a protocol for testing ideas about various mechanisms and their consequences in such systems.

##### Research strategy – Recognizing CD. As CD makes things more difficult, it also makes them more interesting. Although no sound and widely accepted methodology exist to analyze complex systems such as MCs (59), statistical perspective advances (60) promise help, even if it may demand large amounts of data to work effectively. First, high reciprocal CD has to be identified in the system by:

a. Modelling: represent a system in 4D and determine whether one or more dimensions change in response to the state of other dimensions.
b. Comparisons: examine a large sample of similar systems to determine whether some diverge from observed trends without an explanation by the first order CD (Table 2). A lack of explanation might be diagnostic of another order of CD exerting dominant influence.
c. Experiment: track the metrics of 4D while manipulating or using a 'natural' experiment at the micro (lab), meso, or macroscale to determine whether reciprocal, non-linear effects occur.

##### Research strategy – investigating high CD systems. Next, to refine grounds for a theoretical framework and testable hypotheses, two principles for investigating complex systems can help. One relies on reducing a system's complexity, while the other involves searching for the mechanistic explanation at a low level.

- Reducing a system's complexity can occur by abstracting fundamental dimensions (Fig. 2 in the main text) or constraining them by holding some constant (experimentally) or within narrow limits (modeling) and examining outcomes. At the 4D level, it may be best to first pose hypotheses about expected trends in responses variables (species richness, stability, or other ecosystem-level properties), with gradients of species interactions (strength, sign), inter-habitat differences (species independent vs. species dominated), specialization (adaptable vs. restricted habitat use) or dispersal (effectively low vs. high). The answers here will provide coarse, high-variance clouds of cases for which a more specific hypothesis may become feasible using the second strategy.
- Specifically, one can seek the mechanistic explanation at a different resolution – e.g., the second-order interactions (Table 2, main text). Here, when all 4D interact reciprocally, testable predictions examined stepwise may enable uncovering the source of CD. The test result from the first variable can then aid in developing a prediction for the next until the process converges to a solution. A less formal approach might use a combination of educated guesses, available data, and filling in the missing information by simulations.
- Still another possible strategy, in analogy to the sibling problem of treatment interference, may involve designing experiments or data collection to estimate interference itself (CD more broadly) (12) rather than estimating the contribution of individual variables.

It is interesting whether we can test simple hypotheses about the effects, for instance, of the spatial configuration of habitat patches on dispersal and its consequences. In such cases, we probably can if we wish to test whether patch arrangement in space (stochastic, regular, size-structured, close, distant, or any mix of these) affects species richness. Nevertheless, the result will remain suspect until shown otherwise. An acceptable test would demonstrate that the configuration tested relative to a null configuration meaningfully modifies specialization gradient, habitat properties other than those explicitly tested, and species interactions. As in most cases, they will. Various methods hinted at above may help. When the latter is the case, a relatively simple mathematical method called the Walsh-Hadamard transform could be adapted to show how higher-order interactions among species influence a system variable of interest (61).

Testing traditional MC models should best follow the strategies above because CD renders simple tests based on data fitting at best inconclusive, if not unreliable, despite sound statistical results. Although reliance on fundamental dimensions remains firmly at the core of MC, specific tests of various model versions may require novel approaches. Recognition that the 4D not only interact in determining the behavior of a community but also co-determine each other may unlock a sounder, even if the more challenging, path to integration of community processes with broader scale phenomena - an implicit motivation for MC framework. Depending on the level of specificity that an empirical test requires, the dialog between the framework and data will likely be most effective if it proceeds from general to specific predictions. These predictions flow naturally from a hierarchy of interactions (Table 2). The most general predictions will draw on the 4D, while more specific predictions will be nested within general predictions and represent higher-order interactions or modifiers of the 4D predictions. We interpret these interactions as incipient hypotheses of increasing specificity. In the long run, a promising approach to dealing with CD may involve converting dimensions to fields and linking these fields in ways analogous to physics or sociology, e.g., (62, 63).

References (Supplementary)

1.	Deutsch, D., *Constructor theory.* Synthese, 2013. **190**(18): p. 4331-4359.

2.	Pansu, J., et al., *Trophic ecology of large herbivores in a reassembling African ecosystem.* Journal of Ecology, 2019. **107**(3): p. 1355-1376.

3.	Atkins, J.L., et al., *Cascading impacts of large-carnivore extirpation in an African ecosystem.* Science, 2019. **364**(6436): p. 173-177.

4.	Leibold, M.A., et al., *The metacommunity concept: a framework for multi-scale community ecology.* Ecology Letters, 2004. **7**: p. 601-613.

5.	Guichard, F., Y.X. Zhang, and F. Lutscher, *The emergence of phase asynchrony and frequency modulation in metacommunities.* Theoretical Ecology, 2019. **12**(3): p. 329-343.

6.	Guzman, L.M., et al., *Towards a multi-trophic extension of metacommunity ecology.* Ecology Letters, 2019. **22**(1): p. 19-33.

7.	Newman, E.A., et al., *Scaling and Complexity in Landscape Ecology.* Frontiers in Ecology and Evolution, 2019. **7**.

8.	Chase, J.M. and M.A. Leibold, *Ecological niches: Linking classical and contemporary approaches*. Interspecific Interactions, ed. J.N. Thompson. 2003, Chicago: University of Chicago Press. 212.

9.	Thompson, P.L., et al., *A process-based metacommunity framework linking local and regional scale community ecology.* Ecology Letters, 2020. **23**: p. 1314–1329.

10.	MacDougall, A.S., et al., *Context-dependent interactions and the regulation of species richness in freshwater fish.* Nature Communications, 2018. **9**(1): p. 973.

11.	Logue, J.B., et al., *Empirical approaches to metacommunities: a review and comparison with theory.* Trends in Ecology & Evolution, 2011. **26**(9): p. 482-491.

12.	Abrams, P.A., *Habitat choice in predator-prey systems: Spatial instability due to interacting adaptive movements.* American Naturalist, 2007. **169**(5): p. 581-594.

13.	Liao, J.B., D. Bearup, and W.F. Fagan, *The role of omnivory in mediating metacommunity robustness to habitat destruction.* Ecology, 2020. **101**(6).

14.	Ryberg, W.A., K.G. Smith, and J.M. Chase, *Predators alter the scaling of diversity in prey metacommunities.* Oikos, 2012. **121**(12): p. 1995-2000.

15.	Vanschoenwinkel, B., F. Buschke, and L. Brendonck, *Disturbance regime alters the impact of dispersal on alpha and beta diversity in a natural metacommunity.* Ecology, 2013. **94**(11): p. 2547-2557.

16.	Haegeman, B. and M. Loreau, *General relationships between consumer dispersal, resource dispersal and metacommunity diversity.* Ecology Letters, 2014. **17**(2): p. 175-184.

17.	Jabot, F. and J. Bascompte, *Bitrophic interactions shape biodiversity in space.* Proceedings of the National Academy of Sciences of the United States of America, 2012. **109**(12): p. 4521-4526.

18.	Yu, D.W., et al., *Experimental demonstration of species coexistence enabled by dispersal limitation.* Journal of Animal Ecology, 2004. **73**(6): p. 1102-1114.

19.	Gravel, D., et al., *Patch Dynamics, Persistence, and Species Coexistence in Metaecosystems.* American Naturalist, 2010. **176**(3): p. 289-302.

20.	Miller, T.E. and J.M. Kneitel, *Inquiline communities in pitcher plants as a prototypical metacommunity*, in *Metacommunities: spatial dynamics and ecological communities*, M. Holyoak, M.A. Leibold, and R.D. Holt, Editors. 2005, University of Chicago Press: Chicago.

21.	Miller, T.E. and C.P. terHorst, *Testing successional hypotheses of stability, heterogeneity, and diversity in pitcher-plant inquiline communities.* Oecologia, 2012. **170**(1): p. 243-251.

22.	Massol, F., et al., *Linking community and ecosystem dynamics through spatial ecology.* Ecology Letters, 2011. **14**(3): p. 313-323.

23.	Case, T.J., et al., *The community context of species' borders: ecological and evolutionary perspectives.* Oikos, 2005. **108**(1): p. 28-46.

24.	Filotas, E., et al., *Positive interactions and the emergence of community structure in metacommunities.* Journal of Theoretical Biology, 2010. **266**(3): p. 419-429.

25.	Gonzalez, A., et al., *Scaling-up biodiversity-ecosystem functioning research.* Ecology Letters, 2020. **23**(4): p. 757-776.

26.	Cadotte, M.W. and C.M. Tucker, *Should Environmental Filtering be Abandoned?* Trends in Ecology & Evolution, 2017. **32**(6): p. 429-437.

27.	Boulangeat, I., D. Gravel, and W. Thuiller, *Accounting for dispersal and biotic interactions to disentangle the drivers of species distributions and their abundances.* Ecology Letters, 2012. **15**(6): p. 584-593.

28.	Fronhofer, E.A., et al., *Condition-dependent movement and dispersal in experimental metacommunities.* Ecology Letters, 2015. **18**(9): p. 954-963.

29.	Ladyman, J., J. Lambert, and K. Wiesner, *What is a complex system?* European Journal for Philosophy of Science, 2013. **3**(1): p. 33-67.

30.	Desjardins-Proulx, P., T. Poisot, and D. Gravel, *Artificial Intelligence for Ecological and Evolutionary Synthesis.* Frontiers in Ecology and Evolution, 2019. **7**(402).

31.	Kimmel, K., et al., *Causal assumptions and causal inference in ecological experiments.* Trends Ecol.Evol., 2021. **in press**.

32.	Yitbarek, S., et al., *Deconstructing taxa x taxa x environment interactions in the microbiota: A theoretical examination.* bioRxiv, 2021: p. 647156.

33.	Martin, J.L., *What Is Field Theory?* American Journal of Sociology, 2003. **109**(1): p. 1-49.

34.	O'Dwyer, J.P. and J.L. Green, *Field theory for biogeography: a spatially explicit model for predicting patterns of biodiversity.* Ecology Letters, 2010. **13**(1): p. 87-95.
